# Enzyme kinetics model for the coronavirus main protease including dimerization and ligand binding

**DOI:** 10.1101/2025.01.01.631001

**Authors:** Van Ngoc Thuy La, Lulu Kang, David D. L. Minh

## Abstract

The coronavirus main protease (MPro) plays a pivotal role in viral replication and is the target of several antivirals against SARS-CoV-2. In some species, CRCs of MPro enzymatic activity can exhibit biphasic behavior in which low ligand concentrations activate the enzyme whereas higher ones inhibit it. While this behavior has been attributed to ligand-induced dimerization, quantitative enzyme kinetics models have not been fit to it. Here, we develop a kinetic model integrating dimerization and ligand binding. We perform a Bayesian regression to globally fit the model to multiple types of biochemical and biophysical data. The reversible covalent inhibitor GC376 strongly induces dimerization and binds to the dimer with no cooperativity. In contrast, the fluorescent peptide substrate has a minor effect on dimerization but binds to the dimer with positive cooperativity. The biphasic concentration response curve occurs because compared to substrate, the inhibitor accelerates turnover in the opposite catalytic site.

## Introduction

In the past few decades, humanity has faced several serious outbreaks of deadly coronaviruses (CoVs).^1,2^ The Severe Acute Respiratory Syndrome (SARS) CoV pandemic from 2002 to 2004 had 8,422 reported cases and a case fatality ratio of 11%. ^2^ The Middle East Respiratory Syndrome (MERS) CoV, which was first identified in 2012, has a case fatality rate of 34.3%. ^2^ As of November 23, 2024, according to the WHO (https://covid19.who.int/), the global COVID-19 pandemic caused by SARS-CoV-2 has resulted in 776,947,553 cases and 7,076,993 deaths.

The main protease (MPro) plays an important role in the CoV life cycle. After viral entry, the release and uncoating of single-stranded RNA is followed by translation of two large open reading frames near the 5’-end, resulting in the production of polyproteins pp1a and pp1ab.^3–5^ The polyproteins undergo post-translational processing through proteolytic cleavage, generating 16 non-structural proteins. These proteins assemble to form the replicase-transcriptase complex, which is associated with the endoplasmic reticulum membrane and plays a crucial role in genomic RNA replication and subgenomic mRNA synthesis.^3,6^ In particular, this process is regulated by two cysteine proteases, the papain-like protease and chymotrypsin-like protease, the latter also known as 3CLpro or MPro. MPro cleaves polyproteins at 11 specific sites.^7–9^ Many CoV drug discovery efforts have focused on inhibiting MPro, thereby disrupting the proteolytic process and the viral replication cycle.^9–11^

Oligomerization has significant impact on the species-dependent activity of MPro. MPro is much more active as a dimer than as a monomer. ^12,13^ While the substrate binding site and active site of MPro exhibit high conservation among CoVs, ^14^ the dimerization interface of MERS-CoV MPro has some key differences from SARS-CoV and SARS-CoV-2^8,15^ that lead to a weaker binding affinity.^13,16^ In MPro variants with weaker dimerization, biochemical assays have yielded unusual biphasic concentration response curves (CRCs),^16,17^ e.g. Figure 2c. The activity of most enzymes decreases monotonically with the addition of inhibitors. Conversely, with these biphasic CRCs, low inhibitor concentrations increase enzymatic activity (the activation phase), while higher concentrations decrease it (the inhibition phase). In both studies, the authors suggested that the biphasic behavior was due to ligand-induced dimerization; binding of an inhibitor to one catalytic site stabilizes the dimerization interface, promoting dimerization and enhancing enzymatic activity in the opposite site. However, the authors did not fit a quantitative model to these data.

The lack of models to fit biochemical CRCs presents a challenge for assessing the quality of an inhibitor and for the optimization of drug leads. Monotonic CRCs are typically fitted with the Hill equation^18^ and the quality of inhibitors summarized by the half-maximal inhibitory concentration (IC50); the lower the IC50, the more potent the inhibitor. With biphasic CRCs, there are two IC50s and it is unclear how they relate to the relevant potency of the inhibitor. For this reason, Barkan et al. ^19^ excluded MERS-CoV (and also HKU9) from screening panels for a broad-spectrum antiviral.

To address this limitation, we have developed an enzyme kinetics model that incorporates dimerization and ligand binding. As the model uses many parameters, a possible concern is overfitting to data. To alleviate this concern, we performed a global fit of the model to multiple biochemical and biophysical datasets of wild type and a double mutant (E290A/R298A) of SARS-CoV-2 MPro from various experimental modalities.^17^ Nonlinear regression usually does not provide an accurate picture of parameter uncertainty.^20^ For this reason, we used Bayesian regression to incorporate prior information about parameters from independent sources and to provide uncertainty estimates. Many posterior distributions of parameters converged well into reasonable ranges consistent with other studies. We have also analyzed the cooperativity of binding for both the substrate and the inhibitor.

## Methods

### Enzyme kinetics model

Our enzyme kinetics model (Figure 1) predicts product formation rates given initial concentrations of monomer, substrate, and a reversible inhibitor. The model is based on the rapid equilibrium assumption - that the equilibration of protein-containing species and product release is much faster than catalysis. It is applicable to initial velocities after equilibration but prior to appreciable product inhibition. The model accounts for concentrations of eleven species: monomer (M), substrate (S), inhibitor (I), monomer-substrate (MS), monomer-inhibitor (MI), dimer (D), dimer-substrate (DS), dimer-inhibitor (DI), dimer-inhibitor-inhibitor (DII), dimer substrate-substrate (DSS), and dimer-substrate-inhibitor (DSI). We will denote each concentration as c*_X_*, where *X* is the symbol for the species. The initial concentrations will be denoted as c*_Mi_*, c*_Si_*, and c*_Ii_* for the monomer, substrate, and inhibitor, respectively.

**Figure 1:**
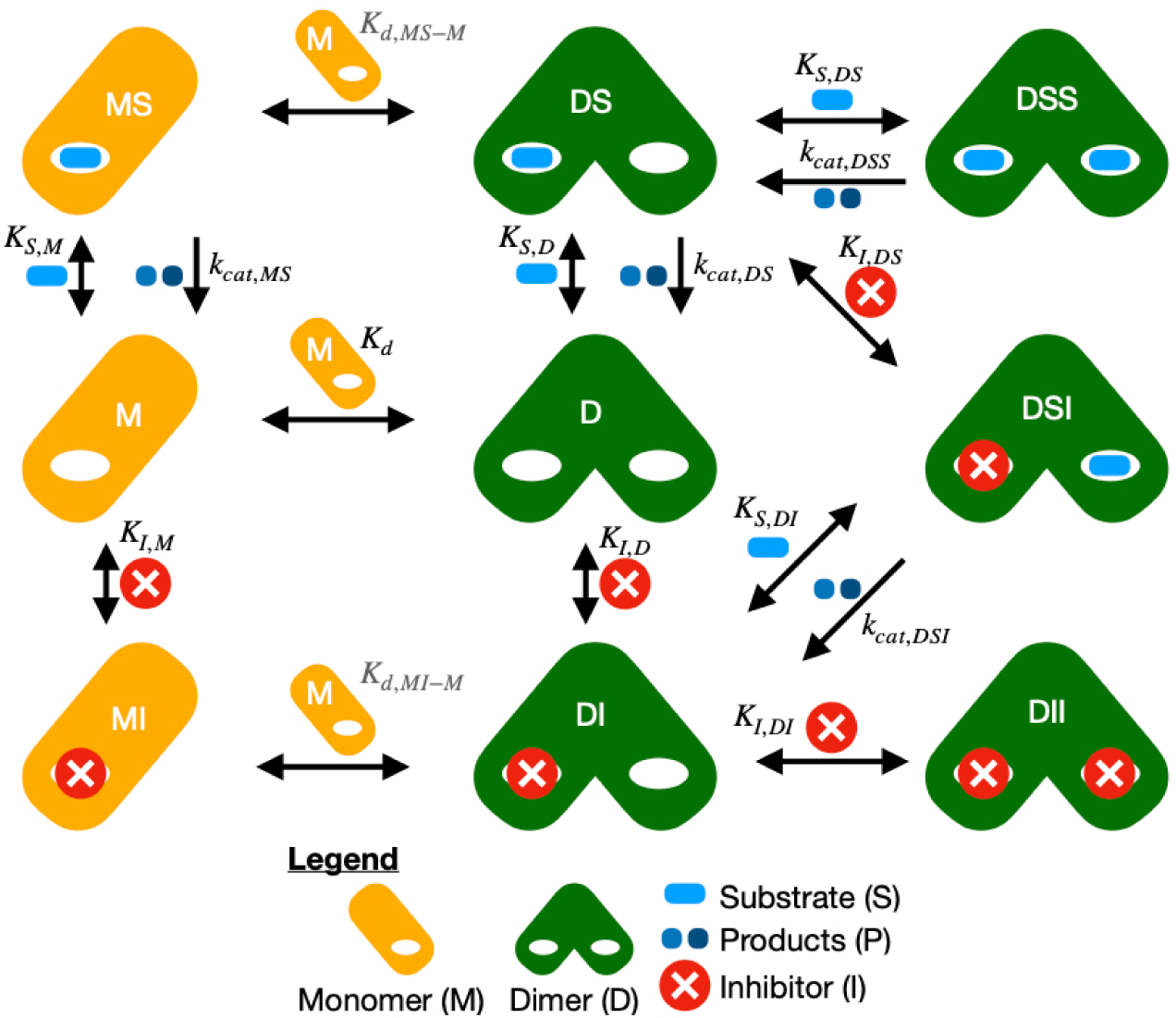
Schematic of the enzyme kinetics model. Proteins are shown as orange rectangles for the monomer (M) or pair of overlapping green rounded rectangles for the dimer (D). Species above the horizontal or tilted arrows are added going right/removed going left. Species to the right of vertical arrows are added going down/removed going up. Equilibrium constants (K) are forward for the direction that leads to more complex species, with K*_d_* for dimerization, K*_I_* for the inhibitor binding, and K*_S_* for the substrate binding. Rate constants k*_cat_* depend on dimerization and ligand binding.

By conservation of mass, the concentrations of these species must add up to the initial concentrations,

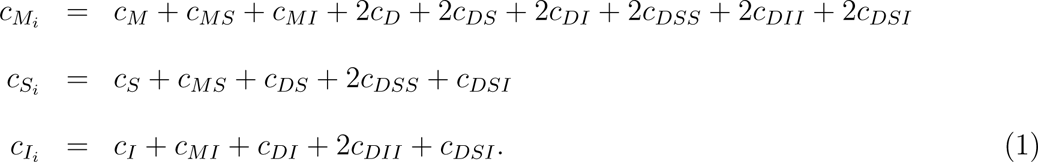

Interconversions between species occur via eleven chemical reactions with dissociation constants,

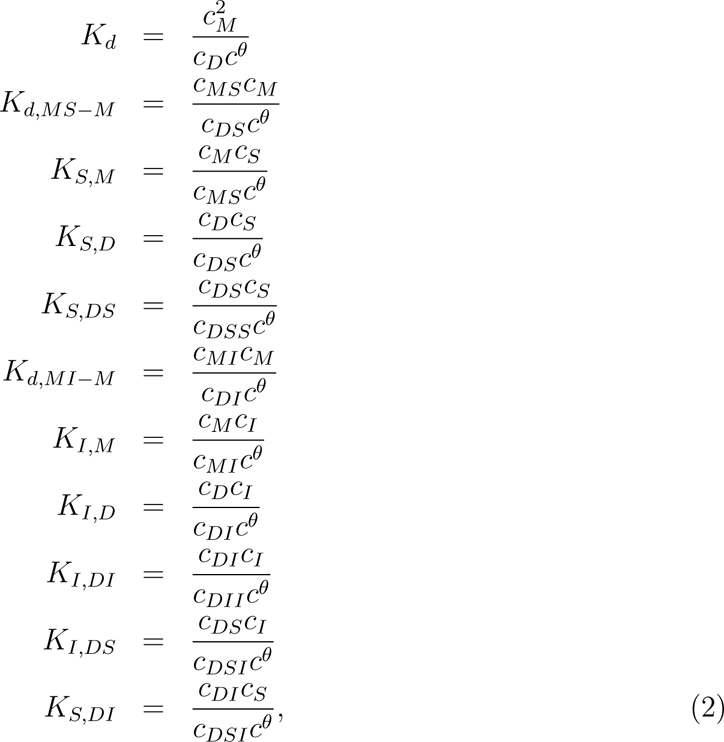

where *c^θ^*= 1 M is the standard concentration.

The equilibrium constants are constrained by three closed loops:

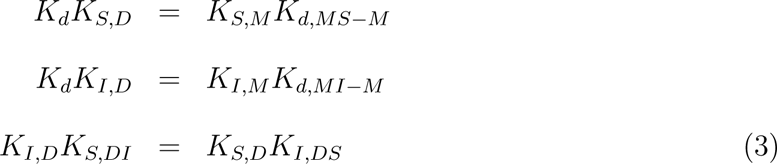

Thus, there are 25 variables - three initial concentrations, eleven concentrations, and eleven equilibrium constants - in 17 equations. If the initial concentrations are specified, the concentrations may be determined based on five equilibrium constants.

In lieu of an analytical solution, we obtained numerical solutions for equilibrium concentrations. We adapted the GeneralBindingModel class (https://github.com/choderalab/assaytools/blob/818d2e57575b4fa6dbf88b45071470698b0c0683/AssayTools/bindingmodels.py#L243) from the open-source Python package *AssayTools*.^21^ GeneralBindingModel implements a function that accepts initial concentrations of different species and a list of mass conservation laws (e.g. Equations 1) and chemical equilibria (e.g. Equations 2) as input and returns equilibrium concentrations as output. For each equation, it defines a target function that is zero when equality is satisfied. To ensure that equilibrium constants and concentrations are positive, the code solves for the logarithm of these quantities. GeneralBindingModel obtains equilibrium concentrations by passing the analytical Jacobian of the target functions to *scipy.optimize.root*.^22^ Key differences between our code (https://github.com/vanngocthuyla/kinetic_mpro/blob/5f61f68a6106bcaad98b55559f93b4d7509fd2d5/scripts/_chemical_reactions.py#L5) and AssayTools are the use of the high performance array computing library *JAX* (https://jax.readthedocs.io/en/latest/) and the Gauss-Newton method, suitable for functional programming in JAX, instead of *scipy.optimize.root*.^22^ Our code also defines three equilibrium constants based on the closed loop constraints (Equations 3).

In addition to using the concentration solver for the full model described here (https://github.com/vanngocthuyla/kinetic_mpro/blob/5f61f68a6106bcaad98b55559f93b4d7509fd2d5/scripts/_kinetics.py#L11), we also used implemented two simplified models: an enzyme-substrate (ES) kinetic model without an inhibitor (https://github.com/vanngocthuyla/kinetic_mpro/blob/5f61f68a6106bcaad98b55559f93b4d7509fd2d5/scripts/_kinetics.py#L87); and a dimer-only model without the monomeric enzyme (https://github.com/vanngocthuyla/kinetic_mpro/blob/5f61f68a6106bcaad98b55559f93b4d7509fd2d5/scripts/_kinetics.py#L404). These models are detailed in Appendix S1 and Appendix S2 of the Supporting Information, respectively. Using simplified models reduces computational cost compared to the full model.

The initial velocity of the enzyme depends on the concentrations and the four rate constants,

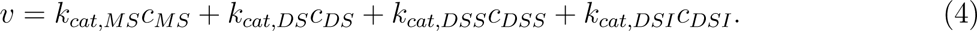

Many biochemical assays for MPro activity measure a response that may be modeled as proportional to the initial velocity, e.g. change in fluorescence signal in a fixed amount of time due to cleavage of a peptide substrate under conditions of excess substrate.^17,23^

### Data

We fit our models to most of the biochemical and biophysical data in a 2022 paper from the John M. Louis group.^17^ The paper includes data for both wild-type MPro and a mutant with a weaker dimerization affinity, denoted as MPro*^Mut^* and MPro*^WT^*. We used data from Figures 3a, 3b, 5b, 5c, 6b, 6d for MPro*^Mut^* of Nashed et al. ^17^ and from Supplementary Figures S2a, S2b, S2c for MPro*^WT^* of the same paper, which was available for download from web site of the journal as Supplementary Data 1. Except for Figure 6b, data were collected from an enzymatic assay in which the substrate and product have different fluorescence resonance energy transfer efficiency. Some assays were performed with the feline coronavirus prodrug GC376, a transition state analog that is a reversible covalent inhibitor of MPro. ^24^ The reported enzyme kinetics parameters^17^ were the velocity of the catalytic reaction (v, M/min) or the catalytic efficiency (K*_m_*/k*_cat_*, min/M). For convenience in model fitting, we converted the catalytic efficiency to its inverse, which we refer to as the Inverse of the Catalytic Efficiency (ICE) (k*_cat_*/K*_m_*, M/min). Data from Figure 6b were total monomer concentrations based on sedimentation velocity experiments performed using analytical ultracentrifugation with enzyme and various concentrations of the inhibitor GC376.

### Bayesian regression

We performed a global fit of the full model to all the data. We inferred the following parameters:

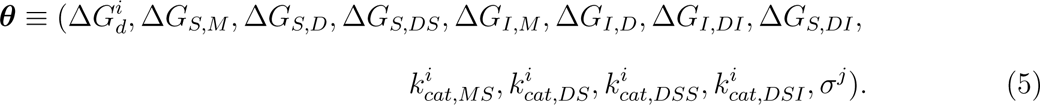

Most of these parameters pertain to the enzyme kinetics model. Free energies Δ*G* are related to equilibrium constants with the same subscripts (Eq. 2) by Δ*G* = −*RT* ln *K*. Dimerization Δ*G_d_* and rate constant parameters *k_cat_* were treated as local to the enzyme variants, in which *i^th^* is the index for MPro*^Mut^* and MPro*^WT^*. The other parameters were treated as global, with the same values in both enzyme variants. In addition to kinetic model parameters, the statistical model includes *σ*, a nuisance parameter for the standard error of measurement per concentration of a species. *σ^j^* is assumed to be constant for all data within a figure from Nashed et al. ^17^, indexed by *j*.

#### Likelihood

For each figure, the data ***D*** ∈ {*y*_1_, *y*_2_, …, *y_n_*} consists of the instrument response for a given concentration. The likelihood function is based on the assumption that each measurement is normally distributed about the model value 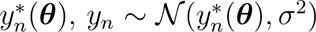. Thus, the likelihood of the data ***D*** is given by,

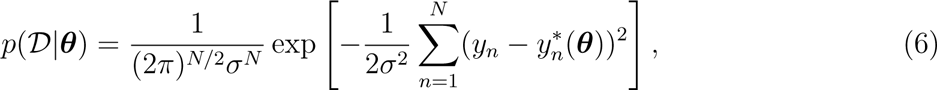

Experimental data were modeled based on concentrations. Reaction rates were modeled based on Equation 4. Catalytic efficiency was modeled by the finite-difference numerical derivative of the reaction rate at *c_S_* = 1 and 2 *µ*M, and ICE by its inverse. Total monomer concentrations in AUC were modeled as the sum of monomer (M) and monomer-inhibitor (MI) concentrations, *c_M,tot_*= *c_M_* + *c_MI_*.

#### Prior

Assuming that the parameters are independent, the prior *p*(***θ***) is a product of the prior for each parameter, *p*(***θ***) = *_i_ p*(*θ_i_*). Based on reported values of dimer-monomer equilibrium of SARS-CoV-2 MPro^25,26^ and binding affinities between the enzyme and the ligand GC376,^17^ normal priors were set for 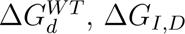 and Δ*G_I,DI_*, Δ*G^W^ ^T^* ∼ Normal(−8.3, 1.2) (kcal/mol)

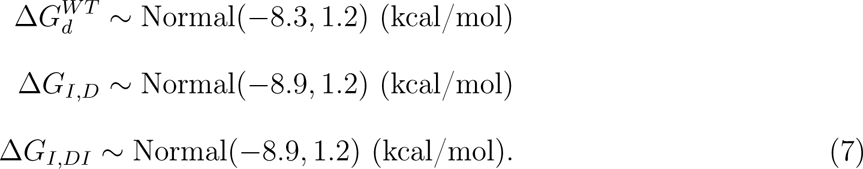

Uniform priors were chosen for 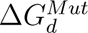 and other binding free energies,

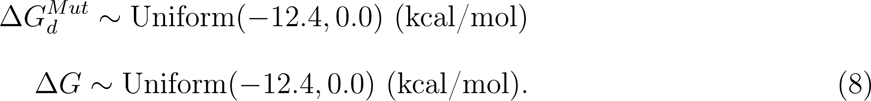

The value of *k_cat,MS_*was set to 0 (min^−1^) for both enzymes based on the assumption that the monomeric state has no enzymatic activity.^13,17^ Uniform priors were set for other rate constants. Because the rate of MPro*^Mut^* was reported as lower than MPro*^WT^*, the upper limit of its prior was set at a lower value,

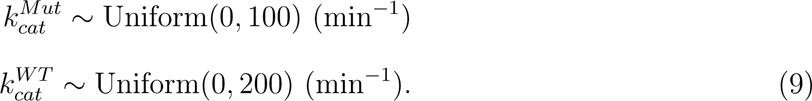

Finally, the uninformative Jeffreys prior was used for *σ*:^27^

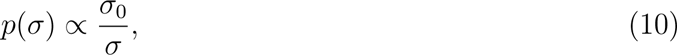

where *σ*_0_ is a reference quantity that simply renders the ratio *σ/σ*_0_ dimensionless. For analytical ultracentrifugation data, *σ*_0_ was set at 1 M; for ICE and kinetic data, *σ*_0_ was set at 1 M/min. The datasets from Figures S2a and S2c were obtained under the same experimental conditions; therefore, we assigned a single *σ* for both datasets.

#### Sampling from the posterior

We employed the No-U-Turn Sampler^28^ to generate Markov Chain Monte Carlo samples from the posterior distributions. ^29^ This sampler automatically determines the number of integration steps for Hamiltonian Monte Carlo. We used the Python implementation in the NumPyro package.^30,31^ After 2,000 warm-up iterations, 10,000 samples were collected for analysis.

All code is freely available at https://github.com/vanngocthuyla/kinetic_mpro/tree/main/sars.

## Results

### Bayesian regression converged and the enzyme kinetics model fit all the data

The convergence of Bayesian posterior sampling was assessed by observing the 5-th, 25-th, 50-th, 75-th, and 95-th percentiles of the marginal probabilities for parameters as the number of Monte Carlo samples increased (Figure S1). With more samples, the percentiles showed minimal variation, accompanied by very small standard errors. This suggests that the posterior distributions were well-explored after only a small number of samples.

The observed parameter set with the highest posterior probability, the Maximum a Posteriori, successfully fit all the data (Figure 2). Trends in the data are reproduced by trends in the model, including the biphasic curve. For Fig. S2a and S2c from Nashed *et al.*,^17^ which were set up under the same experimental conditions, the model appears to be systemically scaled compared to the data (Figures 2g and 2i). This suggests a possible issue in calibration of the catalytic rate in these experiments compared to the others.

**Figure 2:**
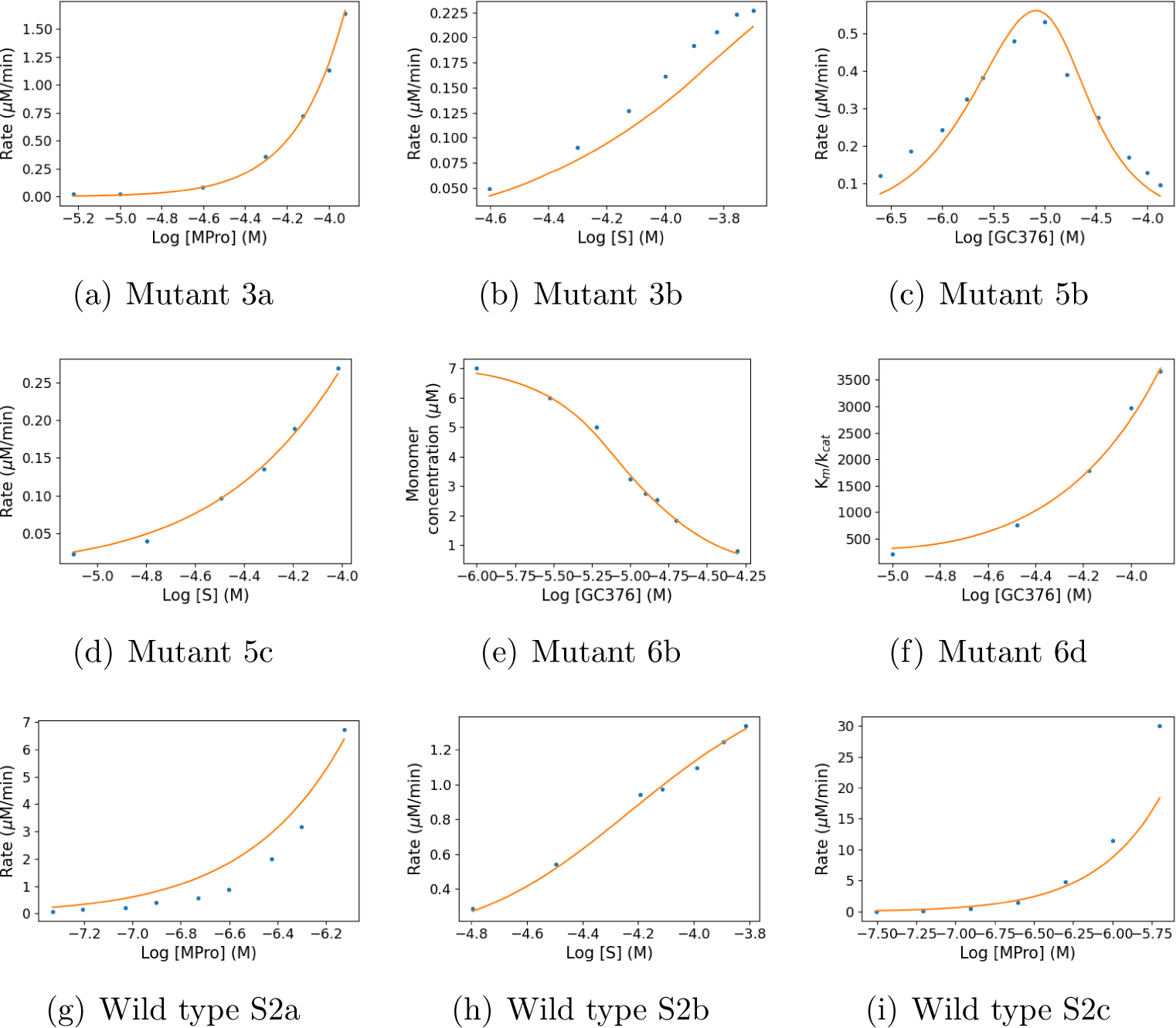
Fit of the model (MAP parameters) to SARS-CoV-2 datasets. X axes are base 10 logarithms of concentrations. The solid line is the theoretical response *y_n_* ∗ (***θ****^MAP^*), where ***θ****^MAP^* is the Maximum a Posteriori estimate of the parameters. Dots are observed responses.

### Many equilibrium constants are tightly determined and some are highly correlated

One-dimensional (1D) marginal distributions of many binding free energies are unimodal, with relatively small highest density intervals (HDIs) (Figure 3). Exceptions include 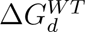, Δ*G_S,DS_*, and Δ*G_I,M_*, which exhibit bimodal and broader distributions. Notably, the Δ*G_I,M_*distribution reaches the upper bound of weak affinity; the inhibitor may have little or no affinity to the monomer.

**Figure 3:**
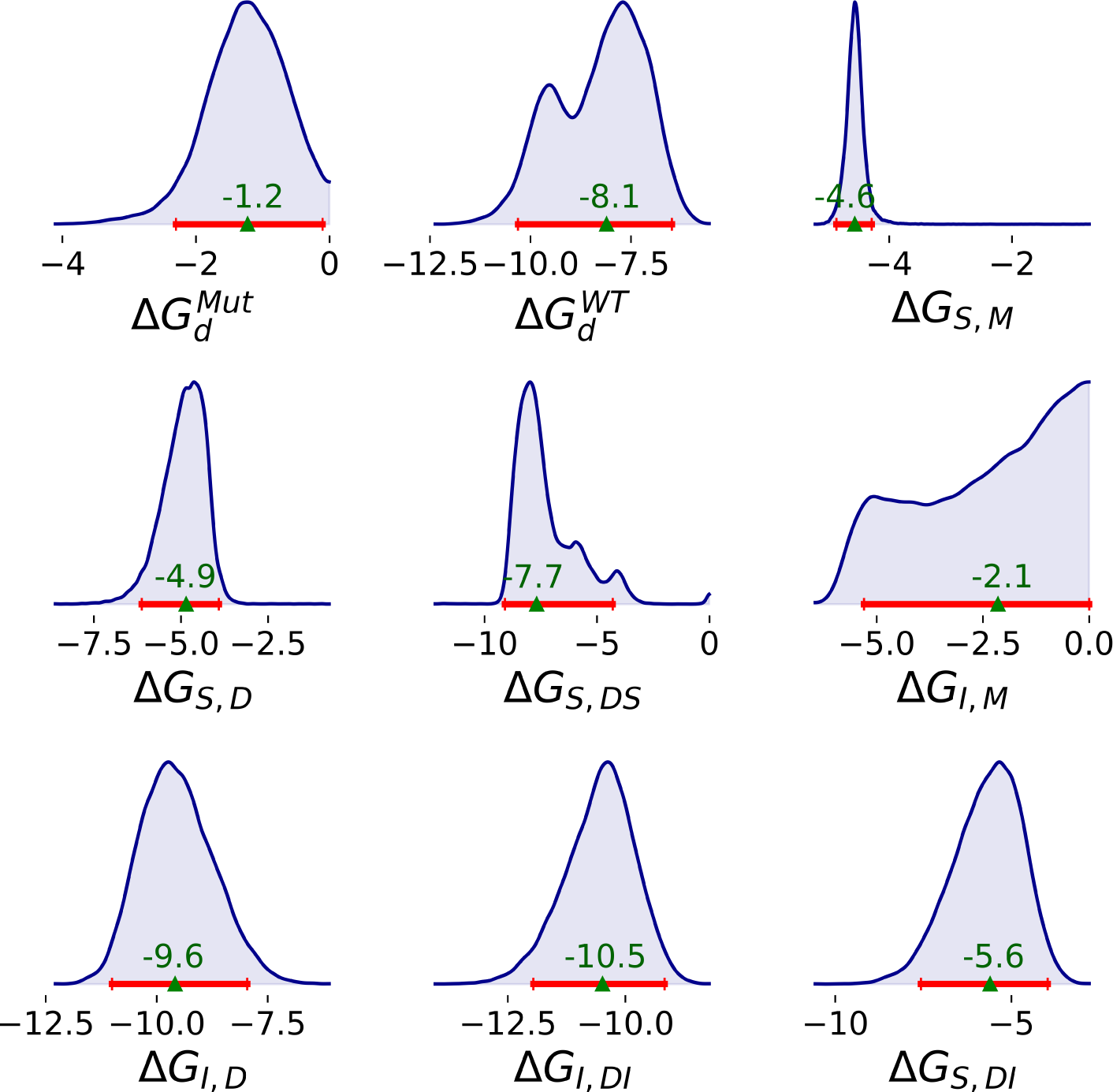
1D marginal distributions for binding free energies (kcal/mol) of SARS-CoV-2 MPro based on 10,000 MCMC samples generated from the Bayesian posteriors. Red line represents 95% HDI. The median of the posterior is shown at the top of the green triangle.

Two-dimensional (2D) marginal distributions reveal that some pairs of parameters exhibit near-independence, while others display strong correlations, with various intermediate levels of correlation observed (Figure 4). Linear correlation was summarized with Pearson correlation coefficients (Figure 5). Δ*G_I,M_*, which has a broad posterior, is uncorrelated with the binding affinities between the inhibitor and the dimeric enzyme, but weakly correlated with several other parameters. There is also minimal correlation between the binding affinities of the substrate and the inhibitor to any form of the enzyme that does not include the other ligand. On the hand, Δ*G_I,DI_* and Δ*G_S,DI_* are positively correlated. Conversely, Δ*G_I,D_* is negatively correlated with both Δ*G_I,DI_* and Δ*G_S,DI_*; the data are well-described by either stronger binding of the inhibitor to the dimer or by weaker binding of the inhibitor and substrate to the dimer-inhibitor complex.

**Figure 4:**
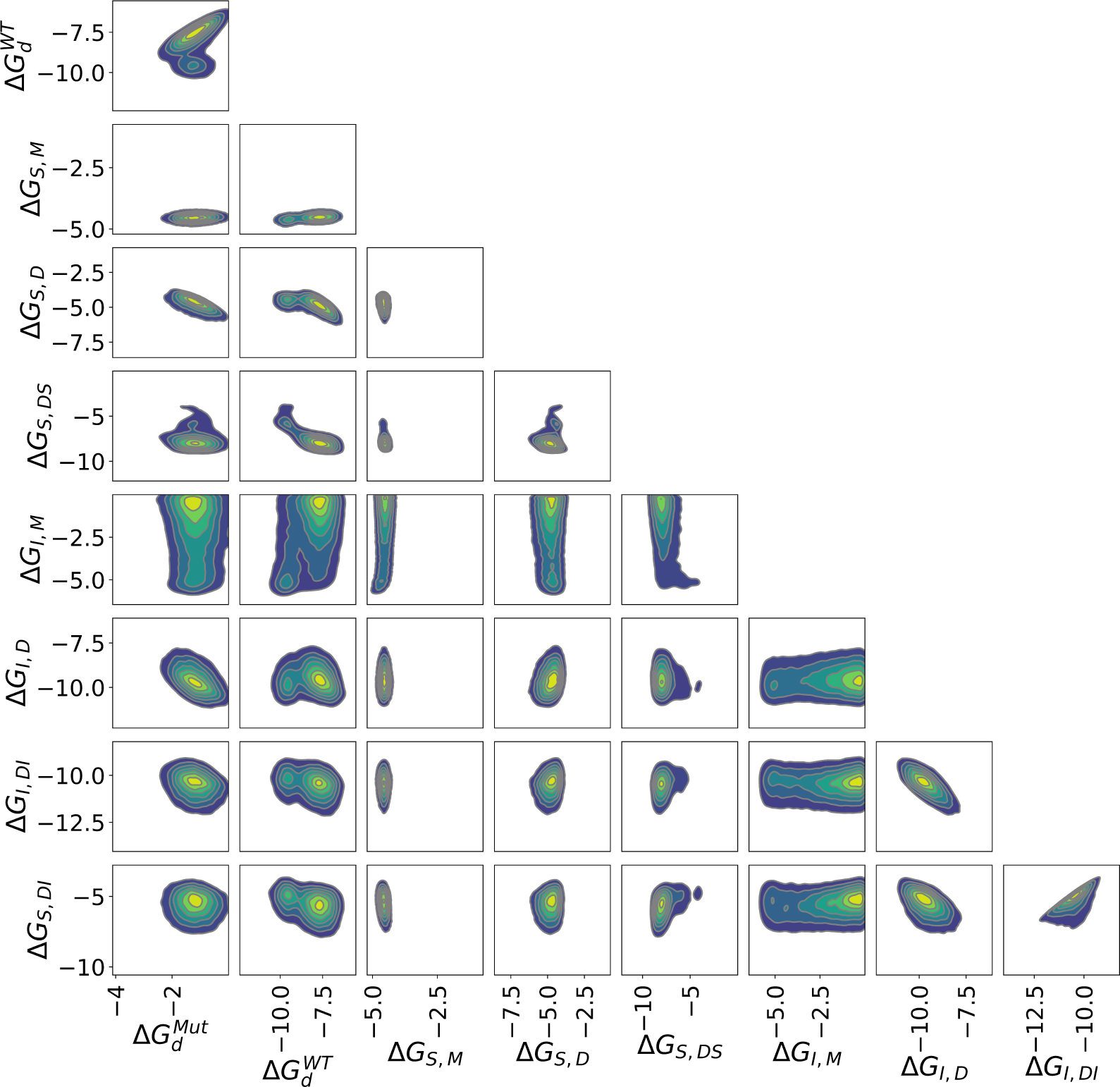
2D joint marginal distributions for pairs of binding free energies (kcal/mol) based on 10,000 MCMC samples from the Bayesian posterior.

**Figure 5:**
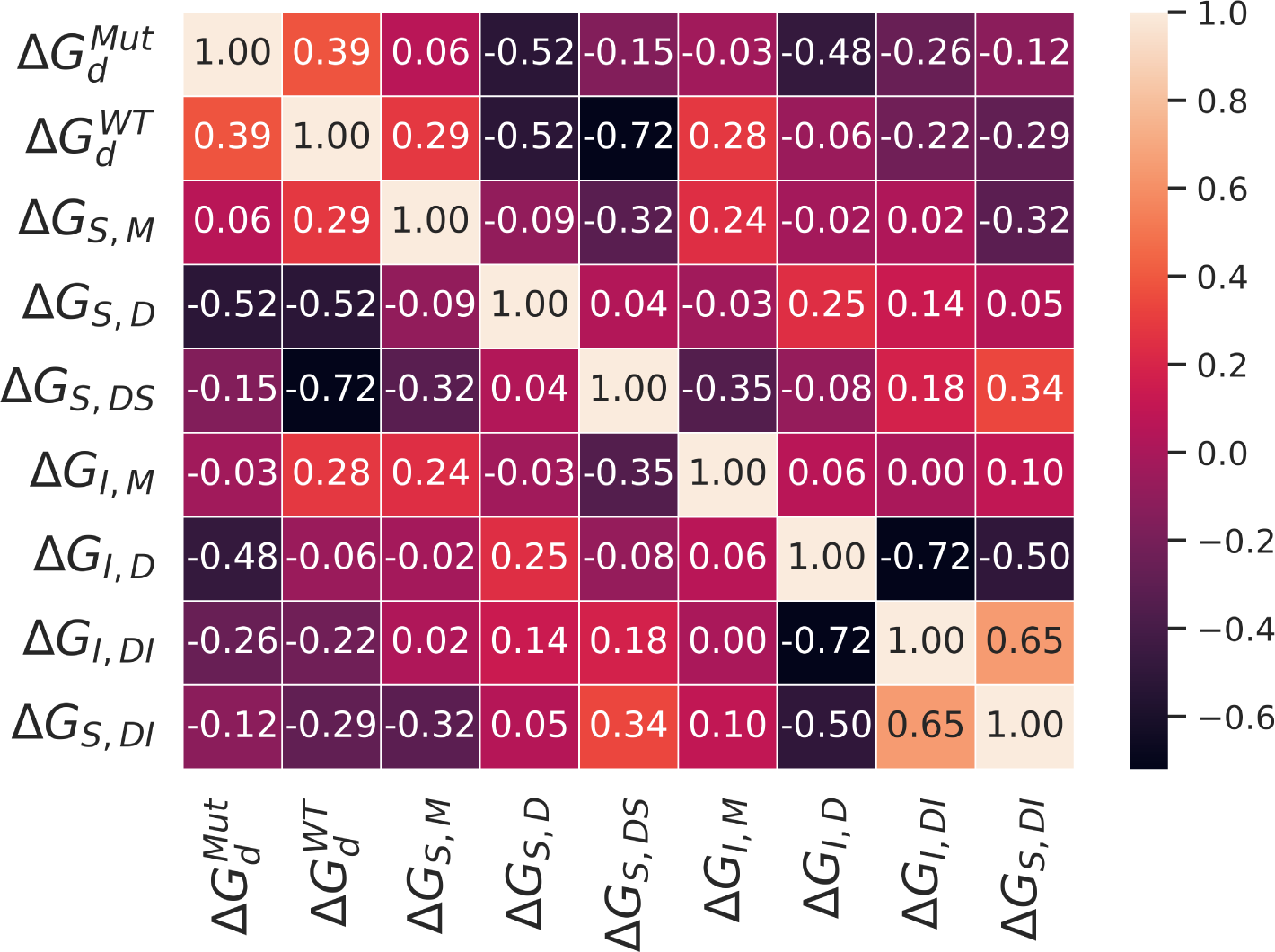
Heat map of the Pearson correlation coefficients estimated from the Bayesian posterior of binding free energies

Besides estimating individual parameter values, we also evaluated differences between binding free energies.

#### MPro dimerization is impacted by mutation and ligand

Mutation and ligand binding have different effects on dimerization. The dimerization free energy of the wild type (median: −8.1; 95% HDI: [−10.3, −6.4] kcal/mol) is much stronger than the mutant (median: −1.3; 95% HDI: [−2.3, −0.1] kcal/mol) (Figure 3). The extent of substrate- or inhibitor-induced dimerization is highly dependent on the ligand. Based on Equation 3, the effects of substrate and inhibitor binding on the dimerization free energy are,

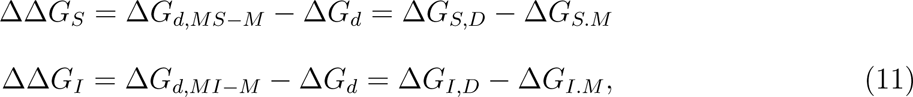

respectively. Binding of GC376 to the monomer strongly favors dimerization (median: −7.1; 95% HDI: [−10.3, −3.7] kcal/mol) (Figure S2). On the other hand, substrate binding has a relatively minor effect on dimerization affinity (median: −0.37; 95% HDI: [−1.7, 0.8] kcal/mol).

#### Ligands exhibit different binding cooperativity

Inhibitor and substrate exhibited distinct binding cooperativity (Figure S3). Although we used the same prior with a mean of −8.9 and a standard deviation of 1.2 kcal/mol for both Δ*G_I,D_* and Δ*G_I,DI_*, the posterior distributions of these two parameters fell into different, though overlapping, ranges. The posterior distribution of the free energy difference between the binding of the first and second inhibitor to the enzyme was broad (Figure S3), indicating no clear evidence of inhibitor-inhibitor cooperativity in the binding of GC376 to MPro. Conversely, while the posterior distributions of Δ*G_S,D_* and Δ*G_S,DS_* also overlap, their free energy difference is larger, suggesting positive cooperativity in substrate binding to the enzyme. Additionally, based on binding free energy differences between the ligands and the dimer-inhibitor complex; the inhibitor binds more tightly to the dimer-inhibitor complex than the substrate.

### The catalytic rate of MPro is affected by mutation and ligand binding

Except for k*_cat,DSS_*, most of the Bayesian posterior distributions of rate constants are broad (Figure S4), suggesting that other data is required for the parameters to be determined. Specifically, we did not include data to estimate *k_cat,DSI_* of the MPro*^WT^*; there were no kinetics measurements of the wild type containing enzyme, substrate, and inhibitor. In addition, the 2D marginal distributions show no correlation between rate constants for either the mutant or wild-type enzyme (Figure S5). Nonetheless, marginal distributions still clearly show that when the enzyme is occupied by two substrates, the catalytic rate of MPro*^WT^* is higher than that of MPro*^Mut^*.

Posterior distributions for ratios of rate constants are more clearly defined than for the rates by themselves (Figure 6). Both the wild type and mutant enzyme show the same trends: catalytic rates of the dimer bound to one substrate (DS) and the dimer-substrate-inhibitor complex (DSI) are approximately equal; and both are much higher than the rate of the dimer bound to two substrates (DSS). The effect is more pronounced for the mutant than the wild type enzyme.

**Figure 6:**
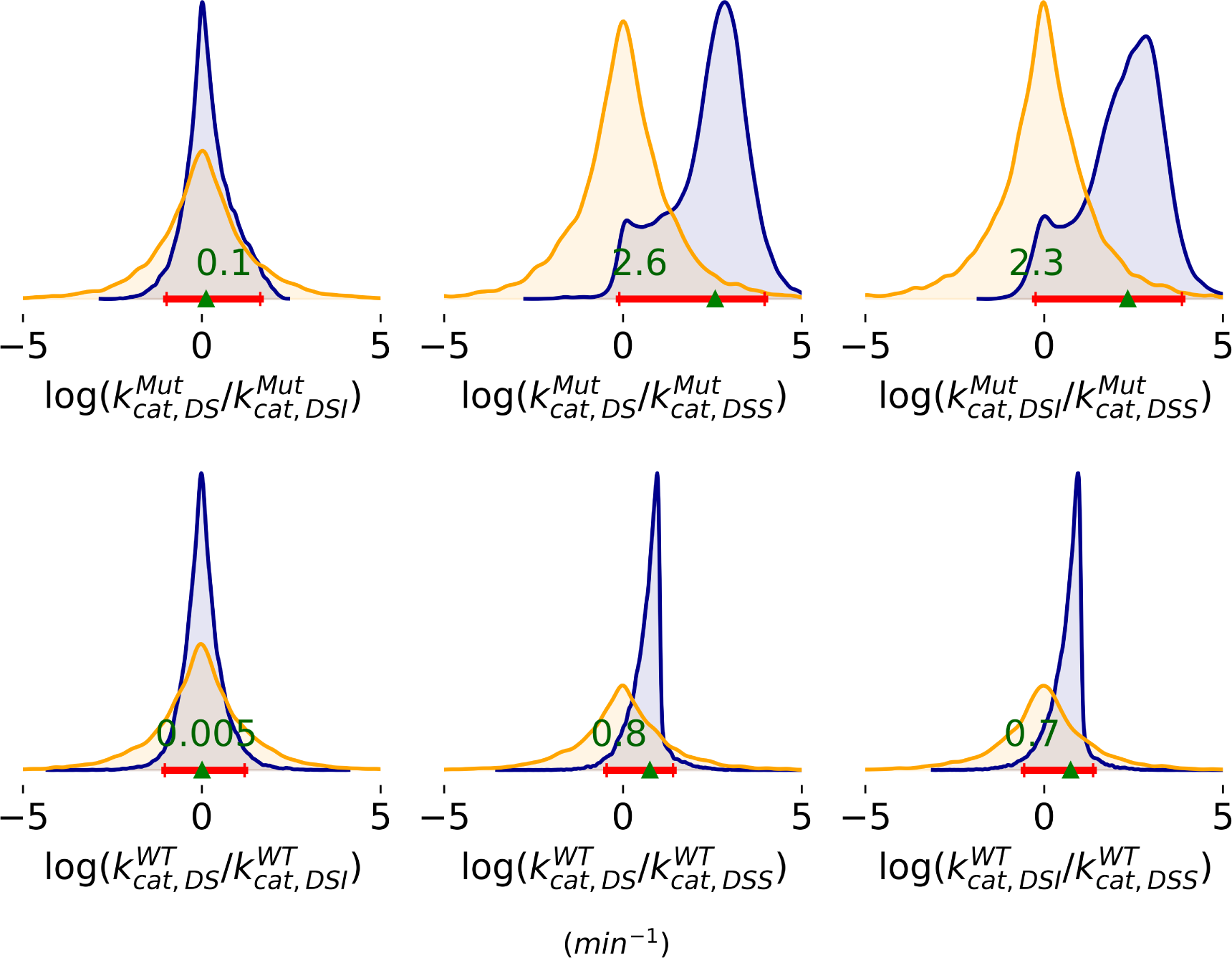
Ratios of rate constants, where the species are DS for dimer bound to a single substrate, DSS for the dimer bound to two substrates, and DSI for the dimer bound to a substrate and inhibitor. Prior distributions are shown in orange and posteriors in blue. Red lines represent 95% HDI. The green triangle marks the median of the posterior.

## Discussion

We developed an enzyme kinetics model that incorporates dimerization and ligand binding and fit it to multiple types of biochemical and biophysical data with SARS-CoV-2 MPro, a substrate, and the inhibitor GC376. The ability of the model to fit to the data suggests that the rapid equilibrium assumption is valid for both variants of this system. By using some informative priors and performing a global fit to data, we were able to achieve narrow Bayesian posterior distributions for many binding free energies. Our analysis also shows that the wild-type enzyme has a much faster catalytic rate than the mutant and that catalytic rates are much faster in the DS and DSI than the DSS state.

By integrating data from multiple experiments, we mitigated the risk of overfitting and achieved more accurate parameter estimation. Data sets contain information about different parameters. For instance, experiments with only enzyme and substrate (e.g. Figures 3a, 3b, 5c, S2a, S2b, and S2c from Nashed *et al.*^17^) provide the most information about enzyme-substrate binding affinities and catalytic rates. Because it contained only enzyme and inhibitor without substrate, Figure 6b of Nashed *et al.*^17^ is especially informative about enzyme-inhibitor binding. While data from Figures 5b and 6d of Nashed *et al.*^17^ involved all three species, these data alone may not achieve narrow posterior distributions for as many parameters as the global fit.

Our work is one of the early examples of applying Bayesian regression to analyzing data from bioanalytical instruments. Most model fitting to scientific data employs nonlinear regression based on maximum likelihood estimation (MLE) of parameters. Unfortunately, MLE usually underestimates parameter uncertainty^20^ due to neglect of entropy.^32^ In contrast, Bayesian regression considers all sets of parameters that enable the model to fit to data. Bayesian regression has been applied to X-ray solution scattering, ^33^ isothermal titration calorimetry,^34–37^ and membrane electrophysiology^38^ data. Bayesian analysis provides more accurate confidence intervals than MLE^34,39^ and insight into which models are best justified by the data.^35,36,38^ Here, we have used Bayesian regression to perform a global analysis of multiple datasets containing different types of data.

Dimerization free energies are consistent with previously reported values of both wild-type and mutant MPro. For wild-type MPro, the 95% HDI of the dimerization free energy of −10.4 and −6.4 kcal/mol is slightly stronger than the value of −6.0 kcal/mol (1.3 *µ*M) reported by Nashed *et al.* but encompasses values of −7.2 kcal/mol (5 *µ*M) based on small-angle X-ray scattering^13^ and −9.4 kcal/mol (0.14 *µ*M) based on mass spectrometry.^40^ In spite of a less informative prior, the posterior for 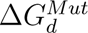 converged into an unimodal distribution, with 95% HDI ranging from −4 to −0.3 kcal/mol, corroborating the weak dimerization of this MPro variant reported by Nashed *et al.*^17^

Our estimates of ligand binding cooperativity are consistent with data from Nashed *et al.*^17^ that were *not* included in our fitting. In addition to the data that we analyzed, Nashed *et al.*^17^ also performed isothermal titration calorimetry of GC376 binding to wild-type and mutant MPro. They found that binding isotherms were well described by a simple binding model with the number of sites *N* close to 1; this ligand binds independently to each subunit. Our finding that GC376 does not have binding cooperativity is consistent with their result. Ligand-induced dimerization affinities are also consistent with previous results. Using small-angle X-ray scattering, the Spinozzi group evaluated how different ligands impact the dimerization affinity of SARS-CoV-2 MPro.^13,26^ Our result that GC376 strongly favors dimerization is consistent with their observation that the inhibitor strengthens the apparent dimerization free energy from −7.2 kcal/mol (5 *µ*M) to −8.7 kcal/mol (0.4 *µ*M). Our observation that the substrate has a minor effect on dimerization (in contrast to GC376) is consistent with their finding that different ligands can have different effects of dimerization free energies. Ligand-induced dimerization has also been observed for SARS-CoV ^41^ and MERS^16^ MPro.

One benefit of our current analysis of ligand-induced dimerization is its broad applicability to other ligand concentrations. The aforementioned analyses ^13,16,26,41^ were based on effective dimerization constants for a specific ligand concentration. In contrast, our rapid equilibrium model may be extrapolated to any ligand concentration. The ability to obtain dimerization parameters based on CRCs of enzyme kinetics may be helpful for the design of dimerization inhibitors, an intriguing mechanism of MPro inhibitors.^42,43^

Our enzyme kinetics model should be readily applicable to other enzyme systems where ligand-induced dimerization has been observed. ^44–47^ One such system is the Kaposi’s sarcoma-associated herpesvirus protease, a weakly associated dimeric serine protease that plays a crucial role in virion maturation.^48^ As in MPro, substrate binding is necessary for dimer stabilization and enzymatic activity^49^ and the extent of dimerization is ligand-dependent.^45,48,49^

According to our model fits to wild-type and mutant SARS-CoV-2 MPro, the primary driver of biphasic CRCs is fast catalytic rate of the mutant DSI state. In the mutant enzyme, the rate constant of DSI is about 200 times higher than that of DSS. The DSI state is most prevalent at intermediate inhibitor concentrations. At low inhibitor concentrations, the less active DS and DSS states are more prevalent. At high concentrations, the inactive state DII dominates. The combination of these factors leads to the biphasic behavior. On the other hand, for the wild-type enzyme, the rate constant of DSI is only about 5 times higher than that of DSS. Because this discrepancy does not exist, CRCs of inhibitors binding to wild-type SARS-CoV-2 MPro have a typical sigmoidal shape.^50^

Along with previous results, our analysis suggests that rate enhancement of the DSI state occurs through allosteric coupling between the dimerization interface and catalytic sites. Nashed et. al. mutated E290 and R298 to alanine. ^17^ Extensive molecular simulations of SARS-CoV-2 show that the transition to an inactive conformation is coupled to the rearrangement of the N terminal residue R4, which in the active conformation forms a salt bridge with E290.^51^ In SARS-CoV MPro, the E290A mutation unsurprisingly has a large effect on both dimerization and catalysis. ^52^ However, R4A weakens dimerization but maintains catalysis.^52^ This finding suggests that it is possible to reshape the dimerization interface in a way that promotes both dimerization and catalysis *in the absence of this salt bridge*. As GC376 binding induces dimerization much more than substrate, it could also provide structural support to the catalytic site to maintain it in an active conformation, accelerating catalysis.

A key contribution of our work is the ability to fit biphasic CRCs of dimeric enzymes to obtain inhibitor binding affinities. Such curves have not only been observed in SARS-CoV-2 MPro with weaker dimerization,^17^ but also in MERS-CoV MPro^16^ and unrelated enzymes such as caspase-1. ^46^ In their analysis of caspase-1 enzyme kinetics, Datta et. al. simulated a biphasic CRC using a steady-state enzyme kinetics model under the assumption that all inhibitor bound irreversibly.^46^ While our model is based on similar states as theirs, there are some important differences between our models and how they were fit to data. Most importantly, our model is suitable for the analysis of reversible noncovalent inhibitors. Another key difference is that our model employs fewer parameters per state; the rapid equilibrium approximation requires only equilibrium constants opposed to forward and reverse rates necessary for the steady-state approximation. A key difference in how the model was fit to data is that we used a Bayesian opposed to standard nonlinear regression. In a subsequent paper under preparation, we plan to describe the application of our model and data fitting procedure to the interpretation of biphasic CRCs for compounds from a drug lead optimization campaign targeting both SARS-CoV-2 and MERS-CoV MPro.

## Conclusion

We developed an enzyme kinetics model that integrates dimerization and ligand binding, suitable for interpreting biochemical data from MPro variants with weak dimerization affinity. Moreover, we demonstrated the use of Bayesian regression for globally fitting multiple datasets, thereby achieving an accurate parameter estimation. Finally, we successfully fitted biphasic CRCs, which have previously been reported without fitting.

## Acknowledgement

We thank John Chodera (ORCID 0000-0003-0542-119X) for helpful discussions and assistance with obtaining funding. Chodera asked us to look into analyzing biphasic CRCs from MERS-CoV MPro. We also thank other members of the AI-driven Structure-enabled Antiviral Platform (ASAP) consortium including Haim Barr and Ed Griffen for encouraging our work and integrating our present analysis into the pan-coronavirus drug lead optimization workflow. This project was supported first by National Science Foundation Award 1905324 to J. Chodera, D. Minh, and L. Kang and then by NIAID of the National Institutes of Health under award number U19AI171399.

This work used the Jetstream2 cloud-based environment at Indiana University through allocation MCB150144 from the Advanced Cyberinfrastructure Coordination Ecosystem: Services & Support (ACCESS) program, which is supported by National Science Foundation grants #2138259, #2138286, #2138307, #2137603, and #2138296.

The content is solely the responsibility of the authors and does not necessarily represent the official views of the National Science Foundation or the National Institutes of Health.

## Supporting Information Available

SI.pdf

includes appendices for two simplified enzymatic kinetic models, figures on the convergence check of Bayesian posteriors, the comparison of dimerization free energy with and without ligand, the comparison of binding free energies, the 1D density distributions of rate constants, the differences between rate constants of MPro, and the 2D distributions for pairs of rate constants.

## Abbreviations Used

CRC: Concentration-Response Curve
CoV: Coronavirus
D: Dimer
DI: Dimer-Inhibitor
DII: Dimer-Inhibitor-Inhibitor
DSI: Dimer-Substrate-Inhibitor
DS: Dimer-Substrate
DSS: Dimer-Substrate-Substrate
IC50: Half-maximal Inhibitory Concentration
HDI: Highest Density Interval
I: Inhibitor
ICE: Inverse of Catalytic Efficiency
MAP: Maximum A Posteriori
MCMC: Markov chain Monte Carlo
MERS: Middle Eastern Respiratory Syndrome
M: Monomer
MI: Monomer-Inhibitor
MS: Monomer-Substrate
MPro: Main Protease
NUTS: No-U-Turn Sampler
SARS: Severe Acute Respiratory Syndrome
S: Substrate
1D: 1-Dimensional
2D: 2-dimensional.

## References

(1) Mirza, A. Z.; Shamshad, H.; Osra, F. A.; Habeebullah, T. M.; Morad, M. An Overview of Viruses Discovered over the Last Decades and Drug Development for the Current Pandemic. European Journal of Pharmacology 2021, 890, 173746.

(2) Bhadoria, P.; Gupta, G.; Agarwal, A. Viral Pandemics in the Past Two Decades: An Overview. Journal of Family Medicine and Primary Care 2021, 10, 2745–2750.

(3) de Wit, E.; van Doremalen, N.; Falzarano, D.; Munster, V. J. SARS and MERS: Recent Insights into Emerging Coronaviruses. Nature Reviews. Microbiology 2016, 14, 523–534.

(4) Guo, Y.-R.; Cao, Q.-D.; Hong, Z.-S.; Tan, Y.-Y.; Chen, S.-D.; Jin, H.-J.; Tan, K.-S.; Wang, D.-Y.; Yan, Y. The Origin, Transmission and Clinical Therapies on Coronavirus Disease 2019 (COVID-19) Outbreak – an Update on the Status. Military Medical Research 2020, 7, 11.

(5) Komatsu, T. S.; Okimoto, N.; Koyama, Y. M.; Hirano, Y.; Morimoto, G.; Ohno, Y.; Taiji, M. Drug Binding Dynamics of the Dimeric SARS-CoV-2 Main Protease, Determined by Molecular Dynamics Simulation. Scientific Reports 2020, 10, 16986.

(6) Gorkhali, R.; Koirala, P.; Rijal, S.; Mainali, A.; Baral, A.; Bhattarai, H. K. Structure and Function of Major SARS-CoV-2 and SARS-CoV Proteins. Bioinformatics and Biology Insights 2021, 15, 11779322211025876.

(7) Hilgenfeld, R. From SARS to MERS: Crystallographic Studies on Coronaviral Proteases Enable Antiviral Drug Design. The Febs Journal 2014, 281, 4085–4096.

(8) Ho, B.-L.; Cheng, S.-C.; Shi, L.; Wang, T.-Y.; Ho, K.-I.; Chou, C.-Y. Critical Assessment of the Important Residues Involved in the Dimerization and Catalysis of MERS Coronavirus Main Protease. PLoS ONE 2015, 10, e0144865.

(9) Lei, J.; Kusov, Y.; Hilgenfeld, R. Nsp3 of Coronaviruses: Structures and Functions of a Large Multi-Domain Protein. Antiviral Research 2018, 149, 58–74.

(10) Osipiuk, J. et al. Structure of Papain-like Protease from SARS-CoV-2 and Its Complexes with Non-Covalent Inhibitors. Nature Communications 2021, 12, 743.

(11) Roe, M. K.; Junod, N. A.; Young, A. R.; Beachboard, D. C.; Stobart, C. C. Targeting Novel Structural and Functional Features of Coronavirus Protease Nsp5 (3CLpro, Mpro) in the Age of COVID-19. The Journal of General Virology 2021, 102, 001558.

(12) Chen, H.; Wei, P.; Huang, C.; Tan, L.; Liu, Y.; Lai, L. Only One Protomer Is Active in the Dimer of SARS 3C-like Proteinase. Journal of Biological Chemistry 2006, 281, 13894–13898.

(13) Paciaroni, A. et al. Stabilization of the Dimeric State of SARS-CoV-2 Main Protease by GC376 and Nirmatrelvir. International Journal of Molecular Sciences 2023, 24, 6062.

(14) Ferreira, J. C.; Fadl, S.; Villanueva, A. J.; Rabeh, W. M. Catalytic Dyad Residues His41 and Cys145 Impact the Catalytic Activity and Overall Conformational Fold of the Main SARS-CoV-2 Protease 3-Chymotrypsin-Like Protease. Frontiers in Chemistry 2021, 9, 692168.

(15) Ferreira, J. C.; Fadl, S.; Rabeh, W. M. Key Dimer Interface Residues Impact the Catalytic Activity of 3CLpro, the Main Protease of SARS-CoV-2. Journal of Biological Chemistry 2022, 298, 102023.

(16) Tomar, S.; Johnston, M. L.; St. John, S. E.; Osswald, H. L.; Nyalapatla, P. R.; Paul, L. N.; Ghosh, A. K.; Denison, M. R.; Mesecar, A. D. Ligand-Induced Dimerization of Middle East Respiratory Syndrome (MERS) Coronavirus Nsp5 Protease (3CLpro). Journal of Biological Chemistry 2015, 290, 19403–19422.

(17) Nashed, N. T.; Aniana, A.; Ghirlando, R.; Chiliveri, S. C.; Louis, J. M. Modulation of the Monomer-Dimer Equilibrium and Catalytic Activity of SARS-CoV-2 Main Protease by a Transition-State Analog Inhibitor. Communications Biology 2022, 5, 160.

(18) Markossian, S. et al., Eds. Assay Guidance Manual [Internet]; Eli Lilly & Company and the National Center for Advancing Translational Sciences: Bethesda (MD), 2004.

(19) Barkan, D. T., et al. Identification of Potent, Broad-Spectrum Coronavirus Main Protease Inhibitors for Pandemic Preparedness. Journal of Medicinal Chemistry 2024,

(20) Kuzmič, P. Methods in Enzymology; Elsevier, 2009; Vol. 467; pp 247–280.

(21) Chodera, J.; Hanson, S.; Isik, M.; Ross, G. A.; Rodríguez-Laureano, L.; Prinz, J.-H. choderalab/assaytools: 0.3.0 experimental release. 2017; 10.5281/zenodo.814975.

(22) Virtanen, P. et al. SciPy 1.0: Fundamental Algorithms for Scientific Computing in Python. Nature Methods 2020, 17, 261–272.

(23) Boby, M. L. et al. Open Science Discovery of Potent Noncovalent SARS-CoV-2 Main Protease Inhibitors. Science 2023, 382, eabo7201.

(24) Arutyunova, E.; Khan, M. B.; Fischer, C.; Lu, J.; Lamer, T.; Vuong, W.; van Belkum, M. J.; McKay, R. T.; Tyrrell, D. L.; Vederas, J. C.; Young, H. S.; Lemieux, M. J. N-Terminal Finger Stabilizes the S1 Pocket for the Reversible Feline Drug GC376 in the SARS-CoV-2 Mpro Dimer. Journal of Molecular Biology 2021, 433, 167003.

(25) Zhang, L.; Lin, D.; Sun, X.; Curth, U.; Drosten, C.; Sauerhering, L.; Becker, S.; Rox, K.; Hilgenfeld, R. Crystal Structure of SARS-CoV-2 Main Protease Provides a Basis for Design of Improved *α*-Ketoamide Inhibitors. Science 2020, 368, 409–412.

(26) Silvestrini, L.; Belhaj, N.; Comez, L.; Gerelli, Y.; Lauria, A.; Libera, V.; Mariani, P.; Marzullo, P.; Ortore, M. G.; Palumbo Piccionello, A.; Petrillo, C.; Savini, L.; Paciaroni, A.; Spinozzi, F. The Dimer-Monomer Equilibrium of SARS-CoV-2 Main Protease Is Affected by Small Molecule Inhibitors. Scientific Reports 2021, 11, 9283.

(27) Jeffreys, H. An Invariant Form for the Prior Probability in Estimation Problems. Proceedings of the Royal Society A: Mathematical, Physical and Engineering Sciences 1946, 186, 453–461.

(28) Hoffman, M. D.; Gelman, A. The No-U-Turn Sampler: Adaptively Setting Path Lengths in Hamiltonian Monte Carlo. 2011.

(29) Neal, R. M. Handbook of Markov Chain Monte Carl; Chapman & Hall/CRC Press, 2012; Vol. 5; pp 113–162.

(30) Bingham, E.; Chen, J. P.; Jankowiak, M.; Obermeyer, F.; Pradhan, N.; Karaletsos, T.; Singh, R.; Szerlip, P.; Horsfall, P.; Goodman, N. D. Pyro: Deep Universal Probabilistic Programming. 2018.

(31) Phan, D.; Pradhan, N.; Jankowiak, M. Composable Effects for Flexible and Accelerated Probabilistic Programming in NumPyro. 2019.

(32) Zuckerman, D. Maximum Likelihood vs. Bayesian Estimation of Uncertainty. 2022.

(33) Minh, D. D. L.; Makowski, L. Wide-Angle X-ray Solution Scattering for Protein-Ligand Binding: Multivariate Curve Resolution with Bayesian Confidence Intervals. Biophysical Journal 2013, 104, 873–883.

(34) Nguyen, T. H.; Rustenburg, A. S.; Krimmer, S. G.; Zhang, H.; Clark, J. D.; Novick, P. A.; Branson, K.; Pande, V. S.; Chodera, J. D.; Minh, D. D. L. Bayesian Analysis of Isothermal Titration Calorimetry for Binding Thermodynamics. PLoS ONE 2018, 13, e0203224.

(35) Nguyen, T. H.; La, V. N. T.; Burke, K.; Minh, D. D. L. Bayesian Regression and Model Selection for Isothermal Titration Calorimetry with Enantiomeric Mixtures. PLoS ONE 2022, 17, e0273656.

(36) Estelle, A. B.; George, A.; Barbar, E. J.; Zuckerman, D. M. Quantifying Cooperative Multisite Binding in the Hub Protein LC8 through Bayesian Inference. PLOS Computational Biology 2023, 19, e1011059.

(37) La, V. N. T.; Nicholson, S.; Haneef, A.; Kang, L.; Minh, D. D. L. Inclusion of Control Data in Fits to Concentration–Response Curves Improves Estimates of Half-Maximal Concentrations. Journal of Medicinal Chemistry 2023, 66, 12751–12761.

(38) George, A.; Zuckerman, D. M. From Average Transient Transporter Currents to Microscopic Mechanism-A Bayesian Analysis. The Journal of Physical Chemistry B 2024, 128, 1830–1842.

(39) La, V. N. T.; Minh, D. D. L. Bayesian Regression Quantifies Uncertainty of Binding Parameters from Isothermal Titration Calorimetry More Accurately Than Error Propagation. International Journal of Molecular Sciences 2023, 24, 15074.

(40) El-Baba, T. J.; Lutomski, C. A.; Kantsadi, A. L.; Malla, T. R.; John, T.; Mikhailov, V.; Bolla, J. R.; Schofield, C. J.; Zitzmann, N.; Vakonakis, I.; Robinson, C. V. Allosteric Inhibition of the SARS-CoV-2 Main Protease: Insights from Mass Spectrometry Based Assays**. Angewandte Chemie International Edition 2020, 59, 23544–23548.

(41) Wei, P.; Li, C.-M.; Zhou, L.; Liu, Y.; Lai, L.-H. Substrate Binding and Homo-Dimerization of SARS 3CL Proteinase Are Mutual Allosteric Effectors. Acta Physico-Chimica Sinica 2010, 26, 1093–1098.

(42) Wei, P.; Fan, K.; Chen, H.; Ma, L.; Huang, C.; Tan, L.; Xi, D.; Li, C.; Liu, Y.; Cao, A.; Lai, L. The N-terminal Octapeptide Acts as a Dimerization Inhibitor of SARS Coronavirus 3C-like Proteinase. Biochemical and Biophysical Research Communications 2006, 339, 865–872.

(43) Goyal, B.; Goyal, D. Targeting the Dimerization of the Main Protease of Coronaviruses: A Potential Broad-Spectrum Therapeutic Strategy. ACS Combinatorial Science 2020, 22, 297–305.

(44) Ching, M. H.; Kaur, P.; Karkaria, C. E.; Steiner, R. F.; Rosen, B. P. Substrate-Induced Dimerization of the ArsA Protein, the Catalytic Component of an Anion-Translocating ATPase. Journal of Biological Chemistry 1991, 266, 2327–2332.

(45) Lazic, A.; Goetz, D. H.; Nomura, A. M.; Marnett, A. B.; Craik, C. S. Substrate Modulation of Enzyme Activity in the Herpesvirus Protease Family. Journal of Molecular Biology 2007, 373, 913–923.

(46) Datta, D.; McClendon, C. L.; Jacobson, M. P.; Wells, J. A. Substrate and Inhibitor-induced Dimerization and Cooperativity in Caspase-1 but Not Caspase-3. The Journal of Biological Chemistry 2013, 288, 9971–9981.

(47) Xu, T.; Gan, Q.; Liu, Q.; Chen, R.; Zhen, X.; Zhang, C.; Liu, J. Substrate-Induced Dimerization of Elaiophylin Glycosyltransferase Reveals a Novel Self-Activating Form of Glycosyltransferase for Symmetric Glycosylation. Acta Crystallographica Section D Structural Biology 2022, 78, 1235–1248.

(48) Shimba, N.; Nomura, A. M.; Marnett, A. B.; Craik, C. S. Herpesvirus Protease Inhibition by Dimer Disruption. Journal of Virology 2004, 78, 6657–6665.

(49) Marnett, A. B.; Nomura, A. M.; Shimba, N.; de Montellano, P. R. O.; Craik, C. S. Communication between the Active Sites and Dimer Interface of a Herpesvirus Protease Revealed by a Transition-State Inhibitor. Proceedings of the National Academy of Sciences of the United States of America 2004, 101, 6870–6875.

(50) Vuong, W.; Khan, M. B.; Fischer, C.; Arutyunova, E.; Lamer, T.; Shields, J.; Saffran, H. A.; McKay, R. T.; van Belkum, M. J.; Joyce, M. A.; Young, H. S.; Tyrrell, D. L.; Vederas, J. C.; Lemieux, M. J. Feline Coronavirus Drug Inhibits the Main Protease of SARS-CoV-2 and Blocks Virus Replication. Nature Communications 2020, 11, 4282.

(51) Nguyen, H. H.; Tufts, J.; Minh, D. D. L. On Inactivation of the Coronavirus Main Protease. Journal of Chemical Information and Modeling 2024, 64, 1644–1656.

(52) Chou, C.-Y.; Chang, H.-C.; Hsu, W.-C.; Lin, T.-Z.; Lin, C.-H.; Chang, G.-G. Quaternary Structure of the Severe Acute Respiratory Syndrome (SARS) Coronavirus Main Protease ^†^. Biochemistry 2004, 43, 14958–14970.

