## Supplementary Information for "Enzyme kinetics model for the coronavirus main protease including dimerization and ligand binding"

### Contents

### Appendix S1: Enzyme-substrate kinetic model without inhibitor

This simplified model does not include any inhibitors. Thus, it excludes species that contain inhibitors and chemical reactions involving the inhibitor may be omitted from the system of equations. The steady-state concentrations of six species - monomer (M), substrate (S), monomer-substrate (MS), dimer (D), dimer-substrate (DS), and dimer-substrate-substrate (DSS) - depend on $c_{M_i}$ and $c_{S_i}$. The steady-state concentrations of these species satisfy five reaction equilibria,

$$\begin{aligned}
K_d &= \frac{c_M^2}{c_D c^\theta} \\
K_{d,MS-M} &= \frac{c_{MS} c_M}{c_{DS} c^\theta} \\
K_{S,M} &= \frac{c_M c_S}{c_{MS} c^\theta} \\
K_{S,D} &= \frac{c_D c_S}{c_{DS} c^\theta} \\
K_{S,DS} &= \frac{c_{DS} c_S}{c_{DSS} c^\theta}
\end{aligned} \tag{1}$$

and there is only one closed loop,

$$K_d K_{S,D} = K_{S,M} K_{d,MS-M}. \tag{2}$$

The two conservation of mass equations are,

$$\begin{aligned}
c_{M_i} &= c_M + c_{MS} + 2c_D + 2c_{DS} + 2c_{DSS} \\
c_{S_i} &= c_S + c_{MS} + c_{DS} + 2c_{DSS}.
\end{aligned} \tag{3}$$

If these initial concentrations are known, then there are eight unknown variables (six concentrations and two equilibrium constants) for the eight equations (five equilibria, one loop, and two conservation). If the equilibrium constants are also specified, the system of equations can be solved for all the steady-state concentrations. The initial velocity of the enzyme is dependent on these concentrations and rate constants,

$$v = k_{cat,MS}c_{MS} + k_{cat,DS}c_{DS} + k_{cat,DSS}c_{DSS}. \tag{4}$$

### Appendix S2: Dimer-only model without monomeric enzyme

If the association constant $K_d$ is large or if there is a high concentration of enzyme, then it can be assumed that there is essentially no monomeric enzyme. Concentrations for species featuring monomer - M, MS, and MI - are zero. Chemical reactions involving the monomer may be omitted from the system of equations. In this case, there are a total of eight species: dimer (D), substrate (S), inhibitor (I), dimer-substrate (DS), dimer-inhibitor (DI), dimer-inhibitor-inhibitor (DII), dimer substrate-substrate (DSS), and dimer-substrate-inhibitor (DSI). The steady-state concentrations of these species satisfy six reaction equilibria,

$$\begin{aligned}
K_{S,D} &= \frac{c_D c_S}{c_{DS} c^\theta} \\
K_{S,DS} &= \frac{c_{DS} c_S}{c_{DSS} c^\theta} \\
K_{I,D} &= \frac{c_{DI} c^\theta}{c_D c_I} \\
K_{I,DI} &= \frac{c_{DII} c^\theta}{c_{DI} c_I} \\
K_{I,DS} &= \frac{c_{DSI} c^\theta}{c_{DS} c_I} \\
K_{S,DI} &= \frac{c_{DSI} c^\theta}{c_{DI} c_S}.
\end{aligned} \tag{5}$$

An additional constraint is introduced by a closed loop,

$$K_{I,D}K_{S,DI} = K_{S,D}K_{I,DS} \tag{6}$$

The three conservation of mass equations are,

$$\begin{aligned} c_{M_i} &= 2c_D + 2c_{DS} + 2c_{DI} + 2c_{DII} + 2c_{DSS} + 2c_{DSI} \\ c_{S_i} &= c_S + c_{DS} + 2c_{DSS} + c_{DSI} \\ c_{I_i} &= c_I + c_{DI} + 2c_{DII} + c_{DSI}. \end{aligned} \tag{7}$$

If the initial concentrations are known, then there are 14 unknown variables (eight concentrations and six equilibrium constants) for the ten equations (six equilibria, one loop, and three conservation). Thus, only four equilibrium constants need to be specified to solve the system of equations. For consistency with the model without inhibitors, it would make sense to choose $K_{S,D}$, $K_{S,DS}$, $K_{I,D}$, $K_{I,DI}$.

The initial velocity of the enzyme is dependent on the steady-state concentrations and the three rate constants,

$$v = k_{cat,DS}c_{DS} + k_{cat,DSS}c_{DSS} + k_{cat,DSI}c_{DSI}. \tag{8}$$

**Figure S1. Convergence of percentiles of the Bayesian posterior for SARS-CoV-2 MPro dataset**

A total of 10,000 samples were obtained from the Bayesian posterior. The results for key parameters are presented, with lines representing the 5-th (blue circle), 25-th (green square), 50-th (red diamond), 75-th (cyan upward triangle), and 95-th (magenta downward triangle) percentiles. Error bars, many too small to be visible, represent the standard deviations calculated from 100 bootstrap samples.

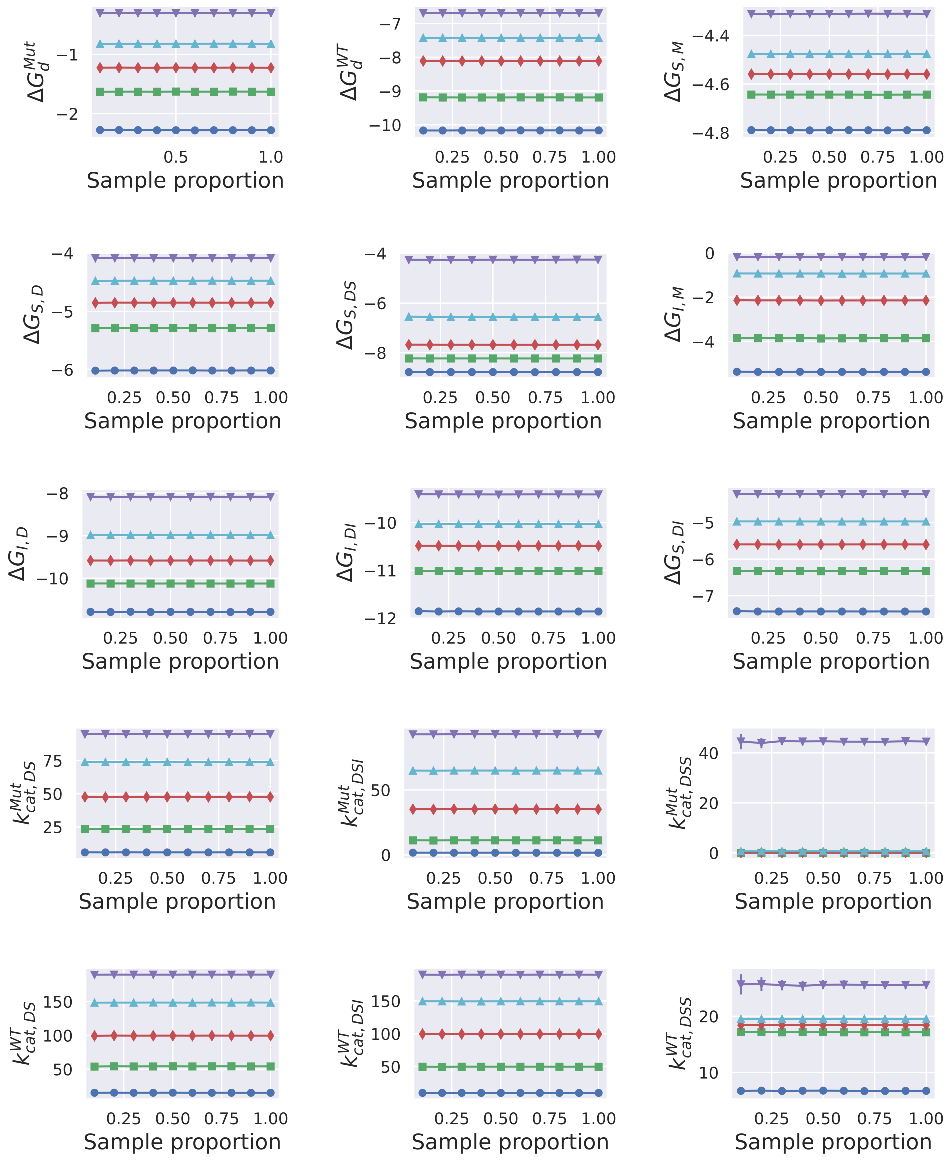

**Figure S2. Differences in dimerization free energies due to ligand binding.**

Red lines represent 95% HDIs. The green triangle marks the median of the posterior.

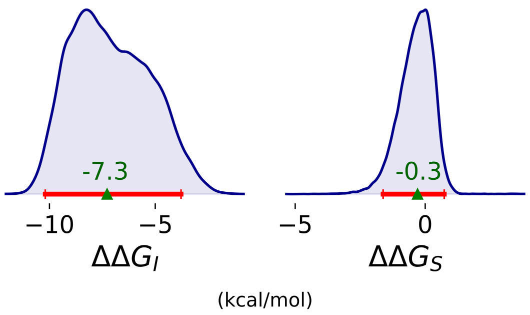


**Figure S3. Differences in binding free energies**

Red lines represent 95% HDI. The green triangle marks the median of the posterior.

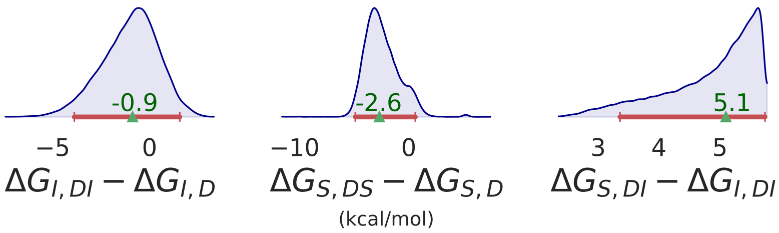

**Figure S4. 1D marginal distributions for rate constants of SARS-CoV-2 MPro**

Distributions were plotted based on 10,000 MCMC samples generated from the Bayesian posterior. Red lines represent 95% HDI. The green triangle marks the median of the posterior. There is some density at the higher rates for $k_{cat,DSS}^{Mut}$ and $k_{cat,DSS}^{WT}$.

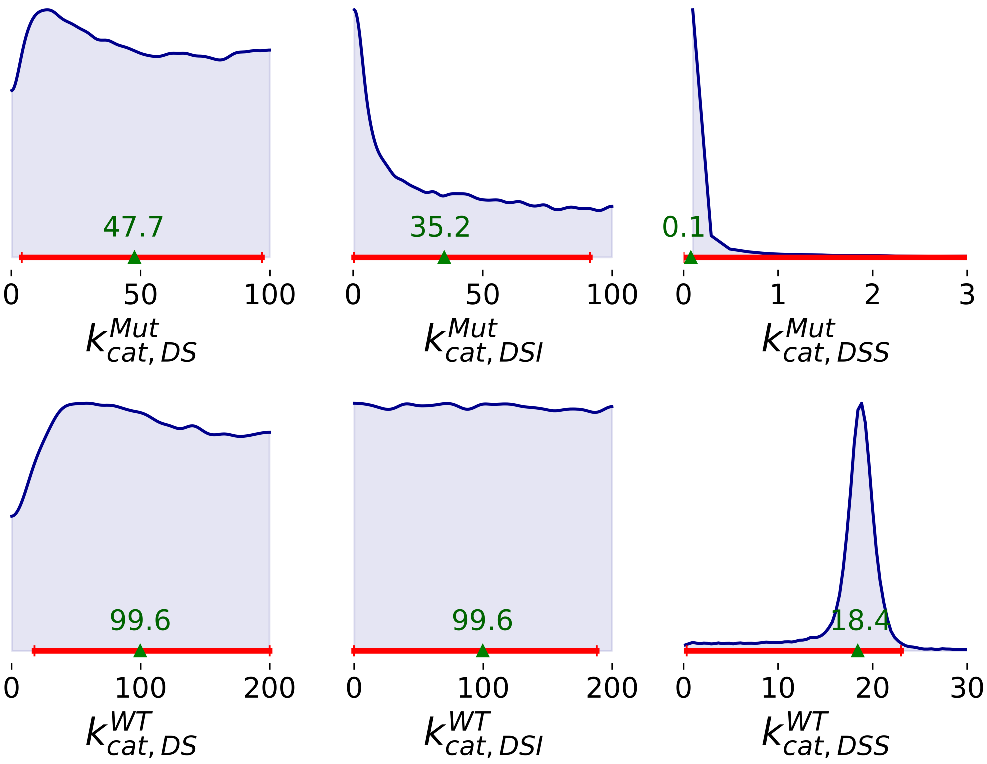

**Figure S5. 2D marginal distributions for pairs of rate constants of SARS-CoV-2 MPro**

2D marginal probability density were plotted based on 10,000 MCMC samples generated from the Bayesian posterior for all datasets.

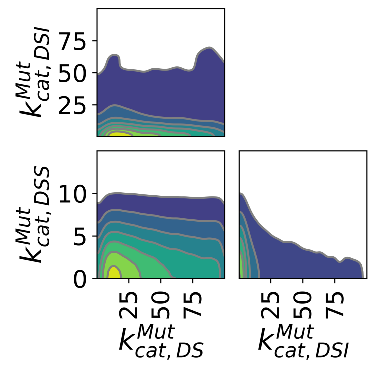


(a) MPro$^{Mut}$

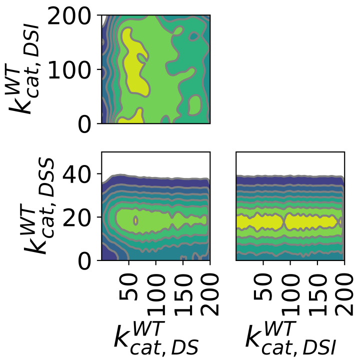


(b) MPro$^{WT}$
